# Auxin is not asymmetrically distributed in initiating Arabidopsis leaves

**DOI:** 10.1101/284554

**Authors:** Neha Bhatia, Marcus G. Heisler

## Abstract

It has been proposed that asymmetric auxin levels within initiating leaves help establish leaf polarity, based in part on observations of the DII auxin sensor. Here we show that the mDII control sensor also exhibits an asymmetry and that according to the ratio-metric auxin sensor R2D2, no obvious asymmetry in auxin exists. Together with other recent findings, our results argue against the importance of auxin asymmetry in establishing leaf polarity.

## Results and Discussion

The leaves of seed plants are usually flat with distinct cell types making up their dorsal (upper) and ventral (lower) tissues. A fundamental question in plant development is how this dorsal-ventral polarity is first specified. Studies based on wounding experiments have suggested the presence of an inductive signal originating from the meristem that promotes dorsal identity in the adaxial (adjacent to the meristem) tissues of the leaf primordium^1,2^. However more recently, it was found that exogenous application of the plant hormone auxin to tomato leaf primordia resulted in the formation of radialized leaves that appeared ventrilized. Hence, relatively high levels of auxin are proposed to promote ventral or inhibit dorsal leaf cell fate^3^. Extending these conclusions, it was also proposed that an asymmetry in the auxin distribution across the leaf adaxial-abaxial axis at leaf initiation acts to help specify leaf polarity during regular development^3^. A critical piece of evidence supporting this latter proposal is that an auxin sensor, the DII^4,5^ indicates low levels of auxin in adaxial leaf tissues compared to abaxial tissues at leaf initiation^3^. Hence asymmetries in auxin concentrations between the adaxial and abaxial leaf tissues, as a result of PIN1 mediated auxin transport, are proposed to help establish leaf polarity^3^. Building on this conclusion, a more recent study proposed that low levels of auxin in adaxial tissues are necessary to restrict the expression of the WOX1 and PRS genes to the middle domain, since auxin promotes their expression^6^. Finally, the reported asymmetry in auxin has also been linked to asymmetries in the mechanical properties of leaf tissues and their morphogenesis^7^.

Despite these studies, reason to reassess the role of any asymmetry in auxin in specifying leaf polarity has arisen due to the recent finding that the expression patterns of genes involved in leaf dorsoventrality are already patterned in meristem tissues where leaves originate, with little change occurring to their expression during organ initiation. Hence leaf polarity appears pre-patterned ^8^. Furthermore, rather than compromise dorsal cell fate, auxin was found to promote dorsal cell fate by maintaining Class III HD-ZIP expression and repressing KAN1 expression in the adaxial cells of organ primordia^8^, which is the opposite of what would be expected according to the auxin asymmetry hypothesis. In addition, Caggiano et al. (2017) also found that exogenous auxin application to young leaf primordia did not influence the spatial pattern of WOX1 and PRS, only the intensity of expression^8^. This also conflicts with the proposal that low levels of auxin are required in adaxial leaf tissues to prevent ectopic WOX1 and PRS expression as previously proposed^6^.

Given the concerns listed above, we decided to examine the distribution of auxin during leaf initiation using not only by the DII sensor assessed in previous studies but also, by looking more closely at the expression pattern of the non-auxin degradable mDII auxin sensor control^4^ as well as the ratiometric R2D2 auxin sensor^9^. We initially focused on the first two leaves three and four days after stratification (DAS), when these leaves are initiating as well as a day later as they begin to elongate and included the PIN1-GFP marker in our analysis to correlate auxin levels with PIN1 expression and polarity. At 3DAS, the expression patterns of REV and KAN1 are already polar within such primordia, although the FIL expression domain at this stage of development is still being refined^8^. Also at this stage, PIN1 is polarized towards the distal tip of leaf primordia but has reversed polarity away from the primordia, towards the meristem, in cells adjacent to the primordia on the adaxial side. According to the ratio-metric auxin sensor R2D2, auxin concentrations were relatively low in adaxial cells of the primordia but also low in abaxial and lateral regions proximally (Fig1-a and d; Fig.S1). High levels of auxin were only found in more distal regions towards the tip of the primordia, matching the overall pattern of signal from PIN1-GFP. No obvious asymmetry in signal between the adaxial and abaxial sides of the primordia was observed. This same overall pattern of signal was found in 16 out of 18 leaves that were examined. At 4DAS, the auxin distribution according to the R2D2 sensor was more uniform although higher levels of auxin appeared associated with the vasculature, again correlating with PIN1 expression (n=12/12 leaves) (Fig. 1 g and j).

**Figure 1.**
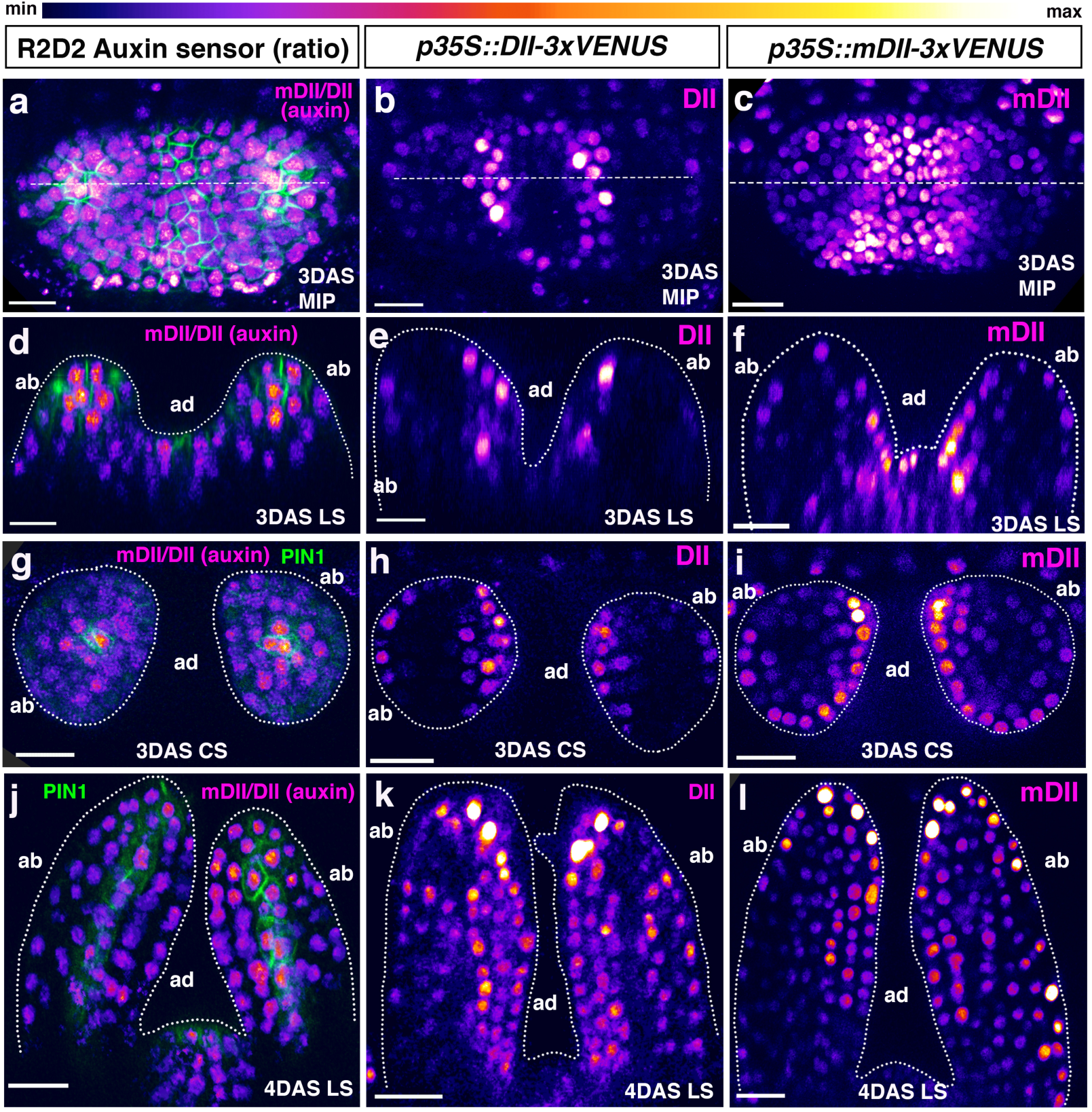
Predicted auxin distribution in young leaves as indicated by different auxin sensors. **(a-c)** Confocal projections of *Arabidopsis* seedlings aged 3DAS (days after stratification) showing predicted auxin distribution based on the ratio-metric auxin sensor R2D2 (magenta) along with PIN1-GFP (green) (a); expression pattern of *p35S::DII-VENUS* (magenta) (b) and *p35S::mutatedDII-VENUS* (mDII, magenta) (c). **(d-f)** Longitudinal reconstructed optical sections of (a-c), respectively, along the dashed lines. **(g-i)** Representative examples of transverse reconstructed optical sections of 3DAS Arabidopsis seedlings showing predicted auxin distribution as indicated by the R2D2 sensor along with PIN1-GFP expression (g), DII-VENUS expression (h) and mDII-VENUS expression (i). R2D2 auxin sensor indicates low but symmetric auxin levels on adaxial and abaxial sides and relatively high auxin levels at the distal tip and in the provasculature where PIN1 expression is also high (d, g). DII-VENUS is more strongly expressed adaxially indicating low auxin levels on the adaxial side of the leaves relative to the abaxial side. However, mDII-VENUS also shows high expression on the adaxial side of the leaf (compare e, h with f, i) and in the shoot meristem. **(j-l)** Representative examples of longitudinal reconstructed optical sections of 4DAS Arabidopsis seedlings showing predicted auxin distribution by R2D2 sensor along with PIN1-GFP expression (green) (i), DII-VENUS expression (j) and mDII-VENUS expression (k). At this stage, the R2D2 sensor indicates a more uniform auxin distribution in the leaf but higher auxin levels in the vasculature where PIN1-GFP expression is also high (j). DII-VENUS shows slightly high expression in adaxial cells and an absence of expression in the vasculature (k). mDII-VENUS shows a similar pattern to DII but is also expressed in the vasculature (l). Scale bars 15µm (a-i) and 20µm (j-l).

As our results using the R2D2 auxin sensor indicate a different auxin distribution compared to that reported previously using the DII marker ^3^, we next re-examined the pattern of DII auxin sensor expression at the same developmental stages. In contrast to the R2D2 pattern, the DII pattern showed an asymmetry of expression in leaf primordia at 3 DAS, indicating relatively low auxin levels in adaxial primordium cells, as found previously. DII signal appeared strongest in the adaxial epidermis but was also stronger in the adaxial sub-epidermal cell layer compared to abaxial epidermal and sub-epidermal cell layers (Fig.1-b, e, h and Fig. S2) (n=18/18 leaves). One day later, although DII signal was still higher in the adaxial cells of the primordia, it started to show a relatively increased expression in the abaxial sub-epidermal layers as well compared to earlier stage (n=13/14 leaves) (Fig1-k). Overall our results using the DII marker are similar to those obtained previously and consistent with the proposal that there are low levels of auxin in the adaxial regions of leaf primordia, in contrast to our results using the R2D2 sensor. Given this discrepancy, we next examined the expression of the mDII sensor which is driven by the same 35S promoter as the DII sensor but is not auxin sensitive. Surprisingly we found that, like the DII results, expression of the mDII marker was also higher in the adaxial cells of the primordia (n= 16/16 leaves) (Fig.1-c,f,i and Fig.S3). The pattern appeared almost identical to the pattern found using the DII marker except that the mDII marker also showed high levels of expression in the shoot meristem whereas the DII sensor did not (compare Fig1-b,c; Fig S2 a-h and Fig S3 a-f). The similarity of expression between DII and mDII was also apparent at 4DAS when the leaves had started to elongate (Fig.1-l) (n=14/14 leaves). To verify the auxin sensitivity of the sensors used we imaged seedlings before and after treatment with 5mM NAA and found a strong decrease in DII expression compared to mDII and an increase in the ratio of VENUS compared to tdTomato signal for the R2D2 sensor, consistent with an increase in auxin levels (Figure S4).

All together these results indicate that the asymmetry in expression found previously for the DII auxin sensor in very young leaf primordia^3^ is not due to an asymmetry in auxin levels but rather, likely due to differences in transcription driven by the 35S promoter used to drive both DII and mDII in adaxial compared to abaxial leaf tissues. We note that although a single section showing control expression of mDII in older leaves was cited by Guan et al., (2017)^10^, our results show that this information was not adequate for assessing similarities and differences between DII and mDII at early developmental stages. To check whether adaxial expression in leaf primordia is a characteristic common to other reporters driven by the 35S promoter, we also examined the expression of 35S::H2B-mRFP1 and 35S::EGFP-LTI6b which had previously been combined into the same plant line by crossing^11^. Surprisingly the expression patterns for these markers were not only different to the mDII marker but also different to each other with the H2B marker showing stronger expression in the adaxial and abaxial epidermis and the LTI6b marker showing a more uniform pattern (Fig. S5). These results reveal that even when the same promoter is utilized to drive FP (fluorescent protein) expression, distinct differences in intensity patterns can occur *in planta*. This may be due to differences in the position of T-DNA insertion in the genome or differences in the surrounding DNA of the vector used. All together then, our results highlight the importance of using ratio-metric sensors for *in vivo* measurements since otherwise it is difficult to adequately control for differences in signal intensity due to promoter activity or other confounding factors. Finally, while the R2D2 marker utilizes a single transgene to express two distinct FPs from two identical promoters^9^, a potentially improved approach to assess auxin levels *in vivo* may be to utilize a single promoter to drive the auxin-dependent degradation domain II from Aux/IAA proteins linked to a tandem fusion of fast and slowly folding FP variants, such as VENUS and mCherry. Such an approach can enable a more direct readout of protein degradation rates^12,13^ and therefore potentially improve measurements of relative auxin concentration.

While our findings are inconsistent with the proposal that asymmetries in the auxin distribution influence leaf polarity^3^, pattern WOX/PRS gene expression ^6^ or regulate tissue mechanics^7^, they also leave other observations unexplained. In particular, to understand the apparent ventrilization of leaf primordia in tomato in response to exogenous auxin^3^ will require further work in assessing how auxin distribution patterns change in response to exogenous treatments and how auxin is distributed during regular development in tomato, preferentially using a ratio-metric auxin sensor such as R2D2.

## Methods

### Plant material and growth conditions

Seeds of the plants expressing *p35S::DII-VENUS* and *p35S::mDII-VENUS* transgenes (*Columbia* ecotype) were obtained from Dr. Teva Vernoux ^4^. An independent batch of seeds expressing *p35S::mDII-VENUS* transgenes (*Columbia* ecotype) was also obtained from NASC (Arabidopsis stock center, NASC ID: N799174) for analysis. R2D2 reporter line carrying *pPIN1::PIN1GFP* transgene (*Landsberg* ecotype) has been described previously^9,14^. Seeds were germinated and grown on GM medium (pH-7 with 1M KOH) containing 1% sucrose, 1X Murashige and Skoog salts (Sigma M5524), MES 2-(MN-morpholino)-ethane sulfonic acid (Sigma M2933), 0.8 % Bacto Agar (Difco), 1% MS vitamins (Sigma M3900) in long day conditions.

### Confocal imaging and data analysis

Seedlings aged 3DAS (days after stratification) and 4DAS were dissected as described previously^14^. Dissected seedlings were then oriented appropriately to obtain a view of the young leaves either from above or from the side. Seedlings were imaged live, on a Leica TCS-SP5 upright confocal laser scanning microscope with hybrid detectors (HyDs) using a 25X water objective (N.A 0.95). VENUS was imaged using argon laser (excitation wavelength 514nm) while tdTomato was imaged using a white light laser (excitation wavelength 561 nm). Z-stacks were acquired in a 512×512 pixel format with section spacing of 1μm and line averaging 2.

Ratio-metric calculations for R2D2 auxin sensor were performed using ImageJ (FIJI, https://fiji.sc) as described previously^14^. All the images were processed using IMARIS 9.0.0 (bit-plane). Optical sections (transverse or longitudinal) were reconstructed using orthogonal slicer in IMARIS.

### Auxin treatment

Seedlings aged 3DAS were dissected imaged and treated with approximately 10μL of 5mM NAA (1-Napthaleneacetic acid) solution in water (0.5M stock in 1M KOH) for 60 minutes and imaged again with same settings as prior to treatment.

## Acknowledgements

We thank Associate Professor Mary Byrne for critical feedback on the manuscript. We thank Dr. Teva Vernoux for kindly providing the seeds of plants expressing *p35S::DII-VENUS* and *p35S::mDII-VENUS* transgenes. We also thank Professor Jim Haseloff for kindly providing the seeds of plants expressing *p35S::H2B-mRFP1* and *p35S::EGFP-LTI6b* transgenes.

---

**FigureS1.**
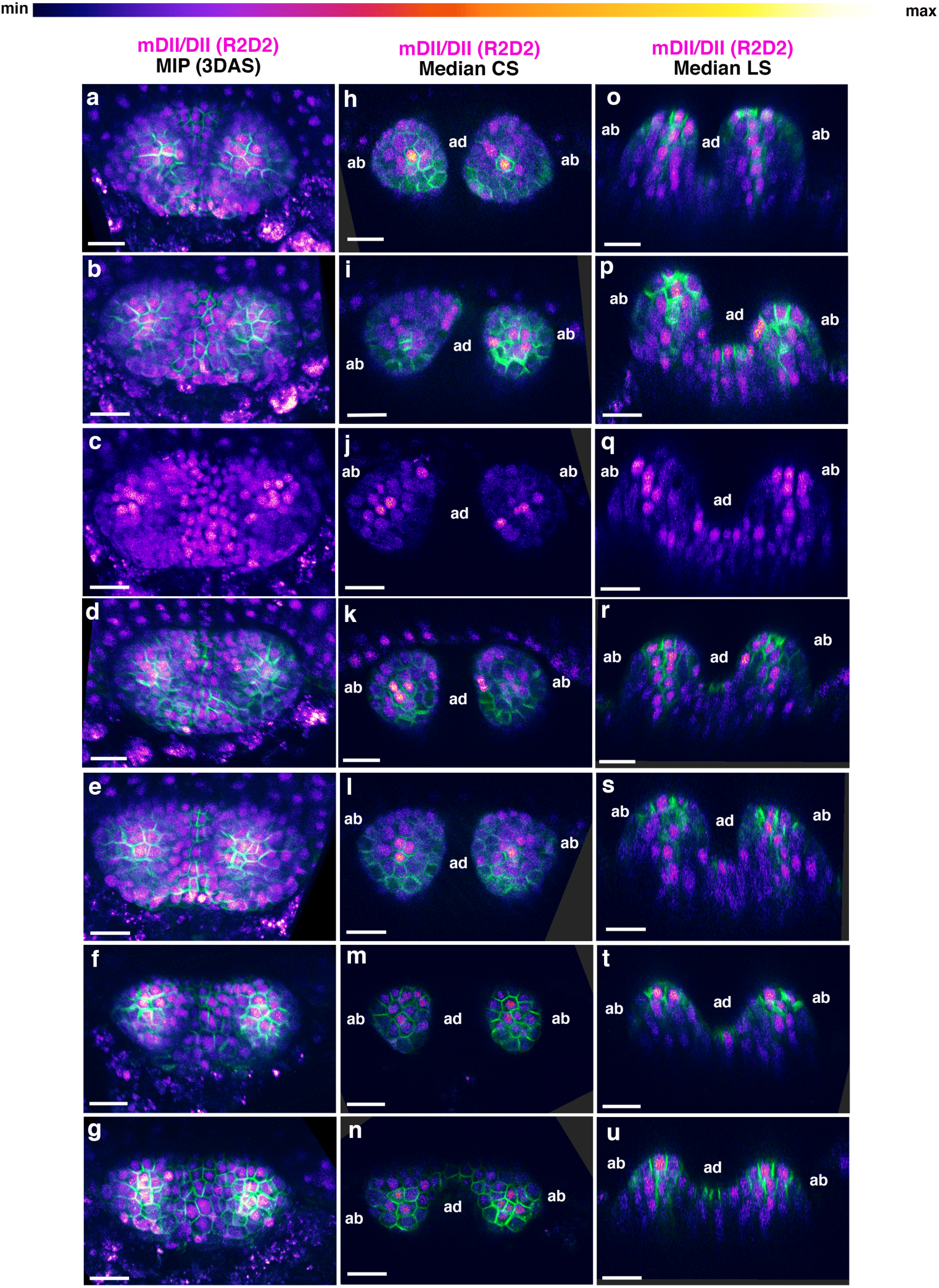
Additional examples of R2D2, ratio-based predictions of auxin distribution in 3DAS old Arabidopsis seedlings. **(a-g)** Confocal projections of *Arabidopsis* seedlings aged 3DAS (days after stratification) showing predicted auxin distribution based on the ratio-metric auxin sensor R2D2 (magenta) along with PIN1-GFP (green). **(h-n)** Median transverse optical sections of (a-g). **(o-u)** Median longitudinal optical sections of (a-g). Scale bars 20µm (a-u).

**Figure S2.**
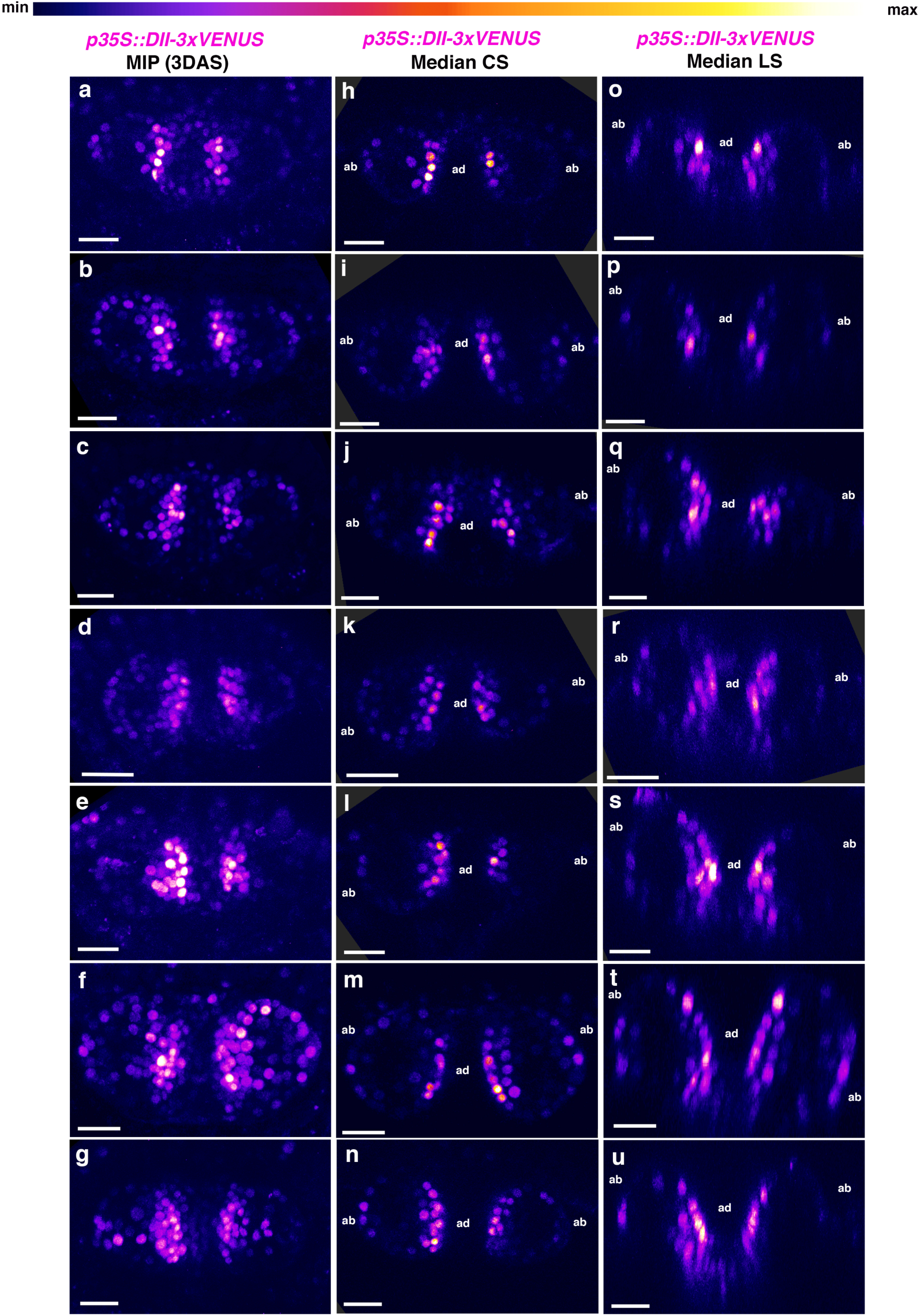
Additional examples *p35S::DII-3xVENUS-N7* expression in 3DAS old Arabidopsis seedlings. **(a-g)** Confocal projections of *Arabidopsis* seedlings aged 3DAS (days after stratification) showing expression pattern of p35s::DII-3xVENUS-N7 sensor (magenta). **(h-n)** Median transverse optical sections of (a-g). **(o-u)** Median longitudinal optical sections of (a-g). Scale bars 20µm (a-u).

**FigureS3.**
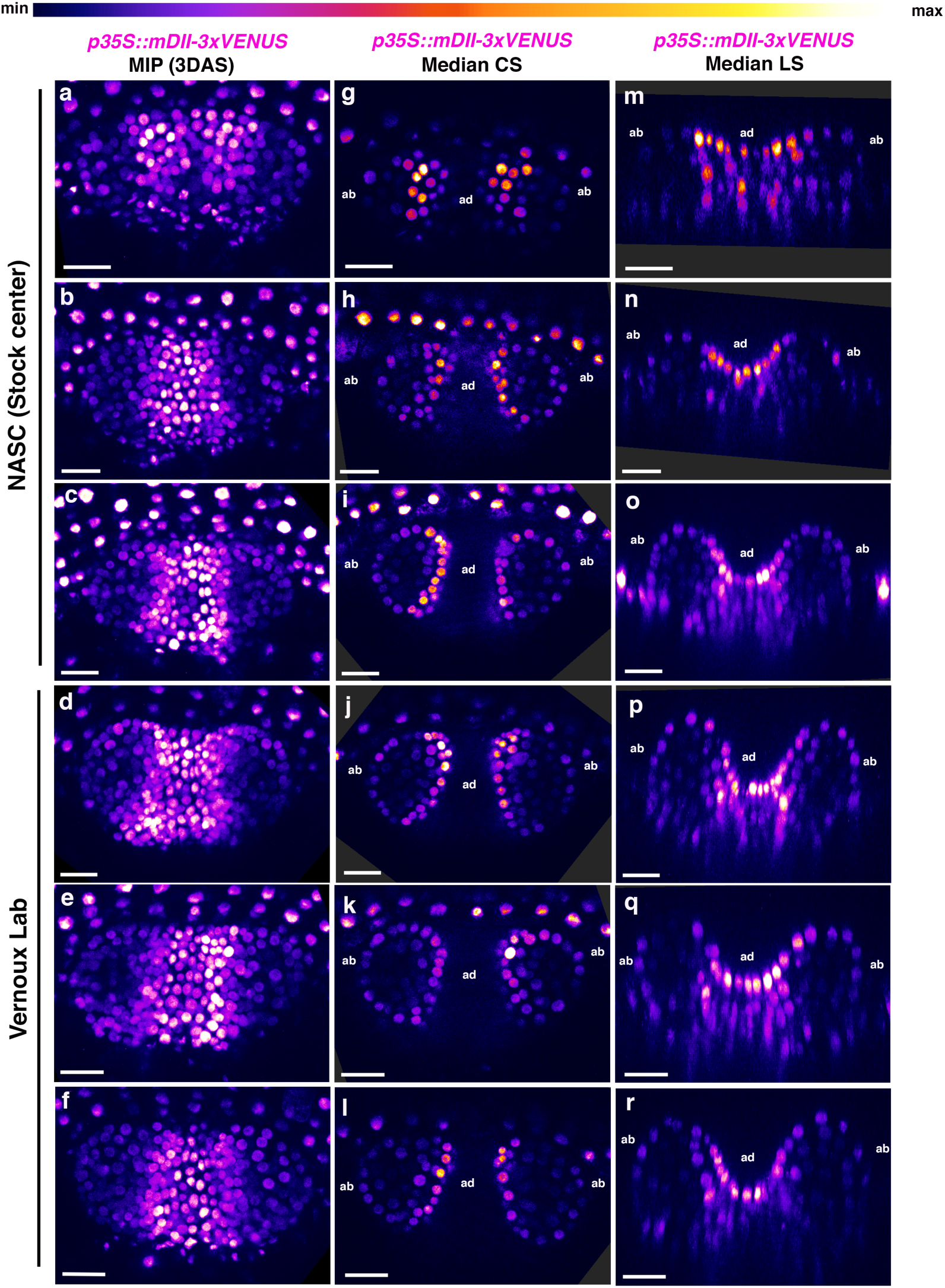
Additional examples of *p35S::mDII-3xVENUS-N7* expression in 3DAS old Arabidopsis seedlings. **(a-f)** Confocal projections of *Arabidopsis* seedlings aged 3DAS (days after stratification) showing expression pattern of p35s::mDII-3xVENUS-N7 sensor (magenta). Note that plants obtained from two different sources showed similar expression pattern. **(h-l)** Median transverse optical sections of (a-f). **(o-r)** Median longitudinal optical sections of (a-g). Scale bars 20µm (a-r).

**FigureS4.**
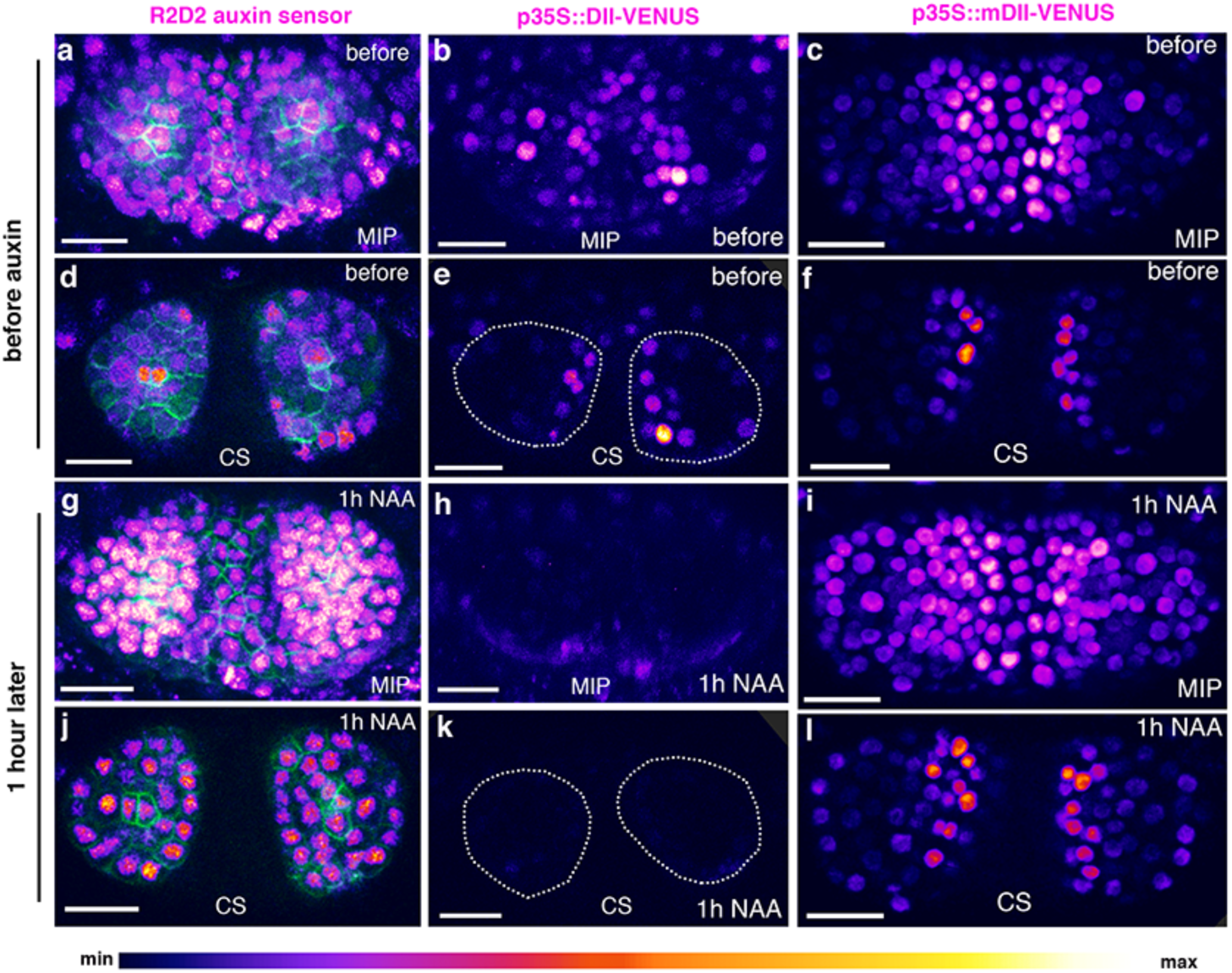
Response of different auxin sensors to external auxin application. **(a-f)** Confocal projections (a-c) and transverse optical reconstructions (d-f) of *Arabidopsis* seedlings, 3DAS, showing predicted auxin distribution based on R2D2 ratio-metric sensor (magenta) along with PIN1-GFP expression in green (a,d), DII-VENUS sensor (magenta) (b,e) and mDII-VENUS sensor (magenta) (c, f) before auxin application. **(g-l)** Confocal projections (g-i) and transverse optical reconstructions (j-l) of *Arabidopsis* seedlings, 3DAS, showing predicted auxin distribution based on R2D2 ratio-metric sensor (magenta) along with PIN1-GFP expression in green (a,d), DII-VENUS sensor (magenta) (b,e) and mDII-VENUS sensor (magenta) (c, f) 1hour after the application of 5mM NAA. Note, R2D2 sensor indicates an increased and broadening of auxin levels after NAA application (compare d and j) (n=3/3). Consistently, DII-VENUS shows an attenuated expression within 1 hour of auxin application (compare b,e with h,j) (n=4/4). mDII-VENUS levels do not decrease upon auxin application (compare c,f with i,l) (n=4/4). Scale bars 20µm (a-l).

**Figure S5.**
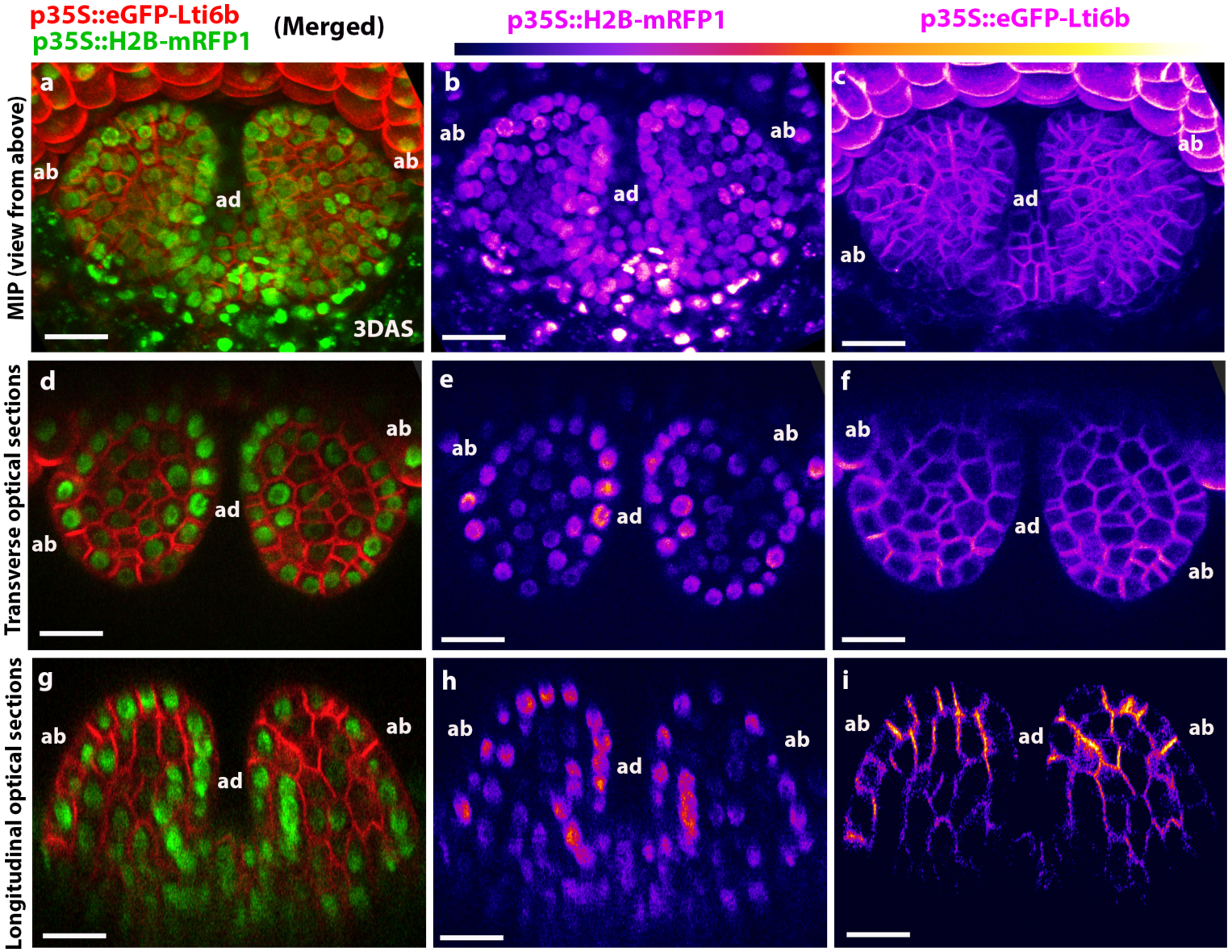
Different 35S promoter-driven reporter genes show different expression patterns in young leaves. **(a-c)** Confocal projections of *Arabidopsis* seedlings aged 3DAS showing expression patterns of a *p35S:eGFP-Lti6b* (membrane marker, green) and *p35S::H2B-mRFP1*(nuclear marker red) together (a), *p35S::H2B-mRFP1* only (magenta) (b) and *p35S:eGFP-Lti6b* only (magenta) (c). **(d-f)** Transverse optical reconstructions of (a-c) respectively; merged (d), *p35S::H2B-mRFP1* only (magenta) (e) and *p35S:eGFP-Lti6b* only (magenta) (f). **(g-i)** Longitudinal optical reconstruction of (a-c) respectively; merged (g), *p35S::H2B-mRFP1* only (magenta) (h) and *p35S:eGFP-Lti6b* only (magenta) (i). Note *p35S::H2B-mRFP1* shows similarly high levels of expression in both adaxial (ad) as well as abaxial (ab) epidermal cell layers and weaker expression in the middle regions (e and h). However, *p35S:eGFP-Lti6b* shows no consistent asymmetry in expression throughout the leaf (f and i). Scale bars 20µm (a-i), (n=10 leaves).

## References

1 Sussex, I. M. Morphogenesis in Solanum tuberosum I: experimental investigation of leaf dorsiventrality and orientation in the juvenile shoot. Phytomorphology 5, 286–300 (1955).

2 Reinhardt, D., Frenz, M., Mandel, T. & Kuhlemeier, C. Microsurgical and laser ablation analysis of leaf positioning and dorsoventral patterning in tomato. Development 132, 15–26 (2005).

3 Qi, J. et al. Auxin depletion from leaf primordia contributes to organ patterning. Proc Natl Acad Sci U S A 111, 18769–18774, doi:10.1073/pnas.1421878112 (2014).

4 Brunoud, G. et al. A novel sensor to map auxin response and distribution at high spatio-temporal resolution. Nature 482, 103–106, doi:10.1038/nature10791 (2012).

5 Vernoux, T. et al. The auxin signalling network translates dynamic input into robust patterning at the shoot apex. Mol Syst Biol 7, 508, doi:10.1038/msb.2011.39 (2011).

6 Guan, C. et al. Spatial Auxin Signaling Controls Leaf Flattening in Arabidopsis. Curr Biol 27, 2940–2950 e2944, doi:10.1016/j.cub.2017.08.042 (2017).

7 Qi, J. et al. Mechanical regulation of organ asymmetry in leaves. Nat Plants 3, 724–733, doi:10.1038/s41477-017-0008-6 (2017).

8 Caggiano, M. P. et al. Cell type boundaries organize plant development. eLife 6, doi:10.7554/eLife.27421 (2017).

9 Liao, C. Y. et al. Reporters for sensitive and quantitative measurement of auxin response. Nat Methods 12, 207–210, 202 p following 210, doi:10.1038/nmeth.3279 (2015).

10 Wang, Y. et al. The Stem Cell Niche in Leaf Axils Is Established by Auxin and Cytokinin in Arabidopsis. Plant Cell 26, 2055–2067, doi:10.1105/tpc.114.123083 (2014).

11 Federici, F., Dupuy, L., Laplaze, L., Heisler, M. & Haseloff, J. Integrated genetic and computation methods for in planta cytometry. Nature Methods 9, 483–U104, doi:10.1038/nmeth.1940 (2012).

12 Khmelinskii, A. et al. Tandem fluorescent protein timers for in vivo analysis of protein dynamics. Nat Biotechnol 30, 708–714, doi:10.1038/nbt.2281 (2012).

13 Barry, J. D., Dona, E., Gilmour, D. & Huber, W. Timer Quant: a modelling approach to tandem fluorescent timer design and data interpretation for measuring protein turnover in embryos. Development 143, 174–179, doi:10.1242/dev.125971 (2016).

14 Bhatia, N. et al. Auxin Acts through MONOPTEROS to Regulate Plant Cell Polarity and Pattern Phyllotaxis. Curr Biol 26, 3202–3208, doi:10.1016/j.cub.2016.09.044 (2016).

